# Methyl Nicotinate Is a Novel Geroprotective Compound That Promotes Mitochondria Dependent Lifespan Extension

**DOI:** 10.1101/2025.05.26.656117

**Authors:** Gao Mingtong, Mohammad Alfatah

**Affiliations:** Healthy Longevity Translational Research Program, Yong Loo Lin School of Medicine, National University of Singapore, Singapore 117544, Singapore; Centre for Healthy Longevity, National University Health System, Singapore 117456, Singapore

**Keywords:** Geroprotective interventions, Methyl nicotinate, lifespan, mitochondria, AMPK, NAD^+^

## Abstract

Aging involves cellular decline and reduced stress resilience. We investigated geroprotective interventions using the yeast chronological aging model and identified methyl nicotinate (MN) as a potent lifespan-extending compound. MN enhanced cellular lifespan and stress resistance through mitochondria-dependent mechanisms, including *AMPK/SNF1* signaling and *HAP4*-mediated mitochondrial biogenesis. These benefits extended to human cells, improving their survival and mitochondrial function under aging conditions. Importantly, the effects of MN are linked to the NAD⁺ biosynthetic pathway, with its conversion to nicotinic acid (NA) and subsequent entry into the NAD⁺ salvage pathway being essential. We also identified the esterase *IAH1* as a key enzyme for converting MN to NA in yeast. Our findings reveal MN as a conserved longevity compound, offering a new avenue for NAD^+^ modulating anti-aging strategies.

## INTRODUCTION

Aging is a complex biological process characterized by a progressive decline in cellular function and stress resilience, ultimately leading to organismal deterioration and age-related diseases ^1–5^. Identifying pharmacological interventions that can delay, or reverse aging processes remains a major goal in biomedical research.

The budding yeast *Saccharomyces cerevisiae* has been instrumental in uncovering fundamental aging pathways due to its genetic tractability and evolutionary conservation ^6–8^. In particular, the chronological lifespan (CLS) model that mimic aging in postmitotic cells, serves as a robust proxy for studying aspects of organismal aging^9–11^.

Several metabolic pathways have been implicated in lifespan regulation, including mitochondrial function, autophagy, and NAD⁺ metabolism. NAD⁺ is a critical cofactor in redox reactions and a substrate for key longevity regulators such as sirtuins and AMPK ^12–19^. Pharmacological enhancement of NAD⁺ levels has been shown to improve mitochondrial function, promote stress resistance, and extend lifespan across species ^14,16,20^. However, most studies have focused on canonical NAD⁺ precursors like nicotinamide riboside (NR) or nicotinic acid (NA) ^21^, while the effects of structurally related compounds remain unexplored.

In this study, we identified methyl nicotinate (MN) as a potent compound that extends cellular lifespan in the yeast chronological aging model. We show that MN delay cellular aging through mitochondria-dependent mechanisms involving *AMPK/SNF1* and *HAP4*-mediated mitochondrial biogenesis. MN also confers resilience against oxidative stress and mild mitochondrial dysfunction, further supporting its role in promoting mitochondrial quality control. We extend these findings to mammalian systems by demonstrating that MN improves survival and mitochondrial function in human HEK293 and SH-SY5Y cells under aging conditions.

Intriguingly, we found that MN interacts with the NAD⁺ biosynthetic pathway. Furthermore, we identified *IAH1*, an esterase gene previously implicated in hydrolysis of short-chain esters, as a key candidate enzyme responsible for converting MN to NA in yeast. Together, our findings uncover a conserved and mechanistically distinct role for MN in promoting longevity and mitochondrial health, with implications for developing novel NAD⁺ modulating geroprotective strategies.

## RESULTS

### Methyl nicotinate extends lifespan during chronological aging in yeast

To identify anti-aging interventions, we conducted a high-throughput chemical screen and identified methyl nicotinate (MN) as a compound that extends the chronological lifespan (CLS) of *Saccharomyces cerevisiae*. We used the prototrophic CEN.PK strain grown in synthetic defined (SD) medium to avoid the confounding effects of amino acid auxotrophy ^22–24^. To validate the screening hit, we performed CLS assays with MN across eight concentrations (0 to 20 mM) using a 2-fold serial dilution. In *S. cerevisiae*, mitotic cells proliferate in glucose-rich conditions and, upon glucose depletion, undergo a diauxic shift to respiratory metabolism, eventually entering a non-dividing stationary phase ^25,26^. These postmitotic cells serve as a model for studying aging in non-dividing human cells ^27,28^.

We first assessed the growth of mitotic cells at 24, 48, and 72 hours and found that MN supplementation did not impair growth, with MN-treated cells showing comparable proliferation to controls (Figure 1A). We then evaluated the effect of MN on CLS using an outgrowth-based survival assay. At various aging time points, cells were transferred to fresh medium, and outgrowth was measured as an indicator of viability. MN significantly extended the lifespan of aging cells (Figure 1B). By day 21, survival reached ∼90% at 5–10 mM MN and nearly 100% at 20 mM, compared to less than 10% survival in untreated cultures (Figure 1B). These results validate our screening outcome and demonstrate that MN robustly extends chronological lifespan in yeast without impairing cell growth under the tested conditions.

**Figure 1.**
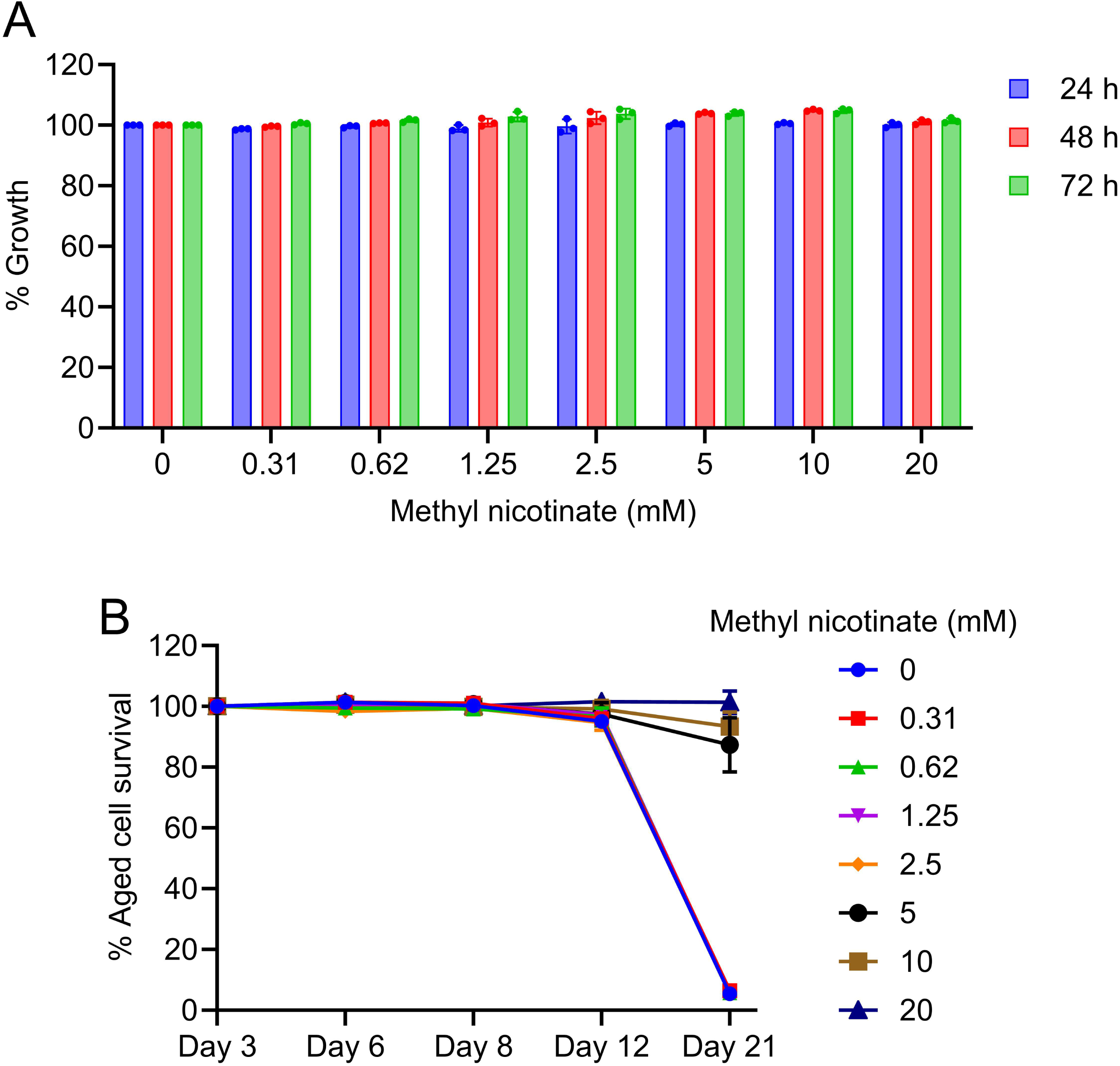
Methyl nicotinate (MN) increases chronological lifespan in *Saccharomyces cerevisiae* prototrophic CEN.PK113-7D cells. (A) Growth analysis of mitotic cells at 24, 48, and 72 hours in synthetic defined (SD) medium with the indicated concentrations of MN in 96-well plate. MN was supplemented in a 2-fold serial dilution ranging from 0 to 20 mM. Data are represented as means ± SD (n=3). (B) Chronological lifespan (CLS) of aging cells treated with the indicated concentrations of MN was assessed in SD medium using a 96-well plate format. Aging Cell survival at various aging time points was measured relative to outgrowth on day 3. Data are presented as mean ± SD (n = 3).

### Methyl nicotinate extends lifespan through AMPK and mitochondrial pathway

Given that MN is a methyl ester of nicotinic acid ^29^, a known NAD⁺ precursor, it may exert its pro-longevity effect by enhancing NAD⁺ metabolism. NAD⁺ is central to mitochondrial function and energy homeostasis and plays a key role in activating sirtuins and *AMPK/SNF1* signaling pathways, both of which are associated with aging and stress resistance ^14–16,18^.

To examine whether AMPK is involved in MN-mediated lifespan extension, we evaluated the growth and chronological lifespan (CLS) of wild type and *AMPK/SNF1*- deleted (*snf1Δ*) cells. MN-treated *snf1Δ* cells showed growth comparable to wild type cells (Figures 2A and 2C), suggesting that MN does not affect cellular growth under these conditions. As expected, *snf1Δ* cells exhibited a shortened lifespan, consistent with the conserved role of AMPK in aging (Figures 2B and 2D) ^30–32^. Interestingly, MN supplementation partially rescued the shortened lifespan of *snf1Δ* cells (Figure 2D), indicating that the anti-aging activity of MN is at least partly mediated through downstream components of the AMPK pathway.

**Figure 2.**
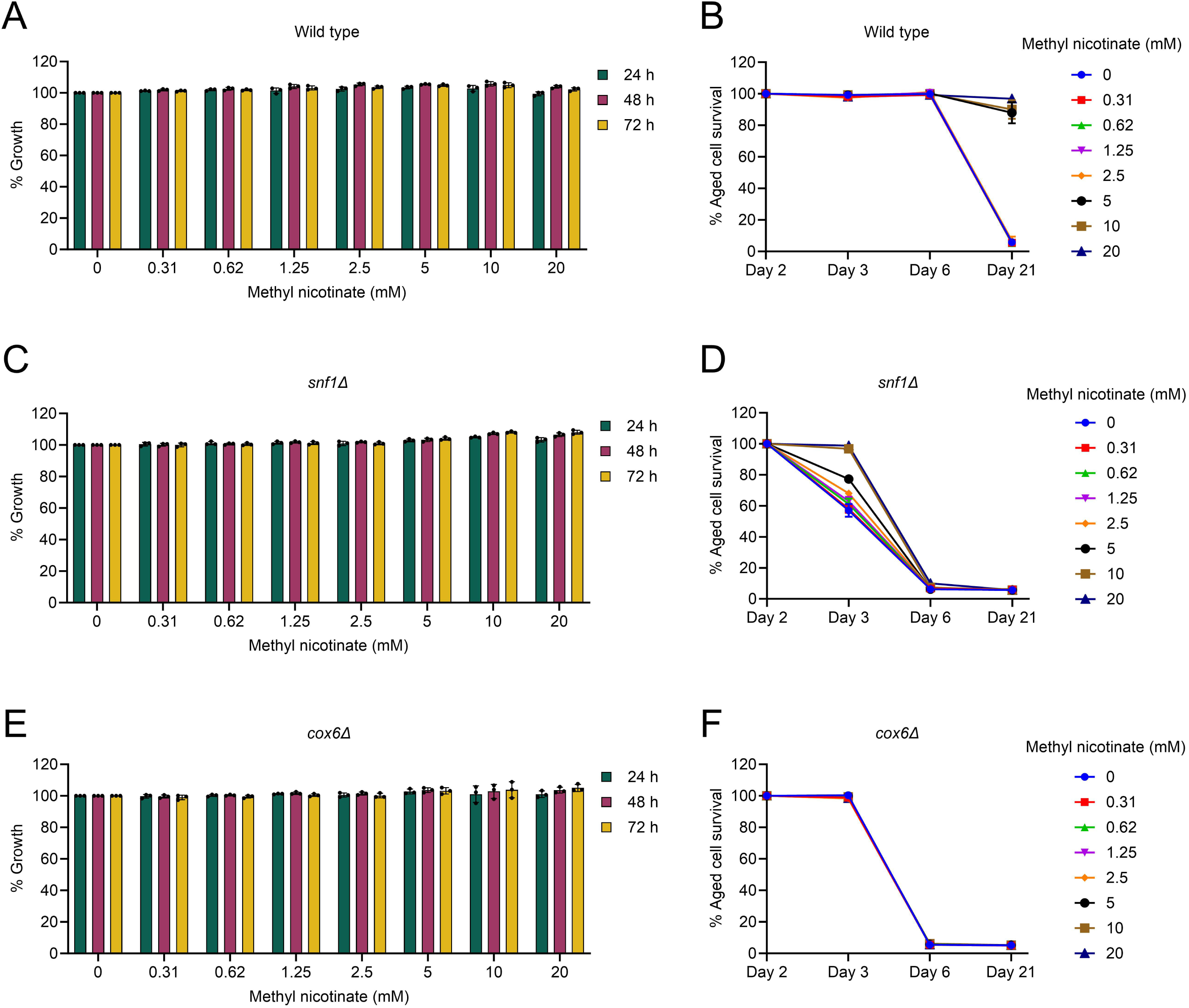
Methyl nicotinate (MN) requires AMPK and mitochondrial function to promote yeast longevity. (A, C, E) Growth analysis of *Saccharomyces cerevisiae* prototrophic CEN.PK113-7D wild type, *snf1Δ*, and *cox6Δ* cells in synthetic defined (SD) medium with indicated concentrations of MN supplementation assessed at 24, 48, and 72 hours using a 96- well plate. Data are presented as mean ± SD (n = 3). (B, D, F) Chronological lifespan (CLS) of wild type, *snf1Δ* and *cox6Δ* cells was evaluated with indicated concentrations of MN supplementation. Survival was measured relative to day 2 outgrowth and plotted over time. Data are presented as mean ± SD (n = 3).

Since NAD⁺ metabolism is also tightly linked to mitochondrial function ^33–35^, we next tested whether mitochondrial function is required for MN’s effects by using the *cox6Δ* mutant, which lacks a subunit of cytochrome c oxidase (complex IV) and exhibits impaired mitochondrial function ^32,36^. Similar to the AMPK-deficient strain, *cox6Δ* cells treated with MN showed growth comparable to wild type cells (Figure 2E). Moreover, *cox6Δ* cells had a markedly shortened lifespan (Figure 2F), consistent with previous reports highlighting the critical role of mitochondria in regulating longevity. Unlike in *snf1Δ* cells, MN supplementation failed to rescue the reduced lifespan of *cox6Δ* cells (Figure 2F), suggesting that functional mitochondria are essential for MN’s geroprotective effect. These findings collectively indicate that MN extends lifespan via a mitochondria-dependent and AMPK-linked mechanism.

### Methyl nicotinate extends lifespan via *SNF1-HAP4*-regulated mitochondrial biogenesis and enhanced quality control mechanisms

Building on these findings, we previously observed that *HAP4* expression—a key regulator of mitochondrial biogenesis—is downregulated in *snf1Δ* cells ^30^, indicating that *SNF1* controls mitochondrial biogenesis through *HAP4*. To further delineate MN’s mechanism, we assessed its effect in *hap4Δ* cells. Growth of *hap4Δ* cells with or without MN was comparable to wild type (Figures 3A and 3C), while lifespan was shortened (Figures 3B and 3D), consistent with prior reports ^32,37^. Notably, MN supplementation completely failed to extend lifespan in *hap4Δ* cells (Figure 3D), reinforcing that mitochondrial biogenesis via *HAP4* is essential for MN’s longevity benefits.

**Figure 3.**
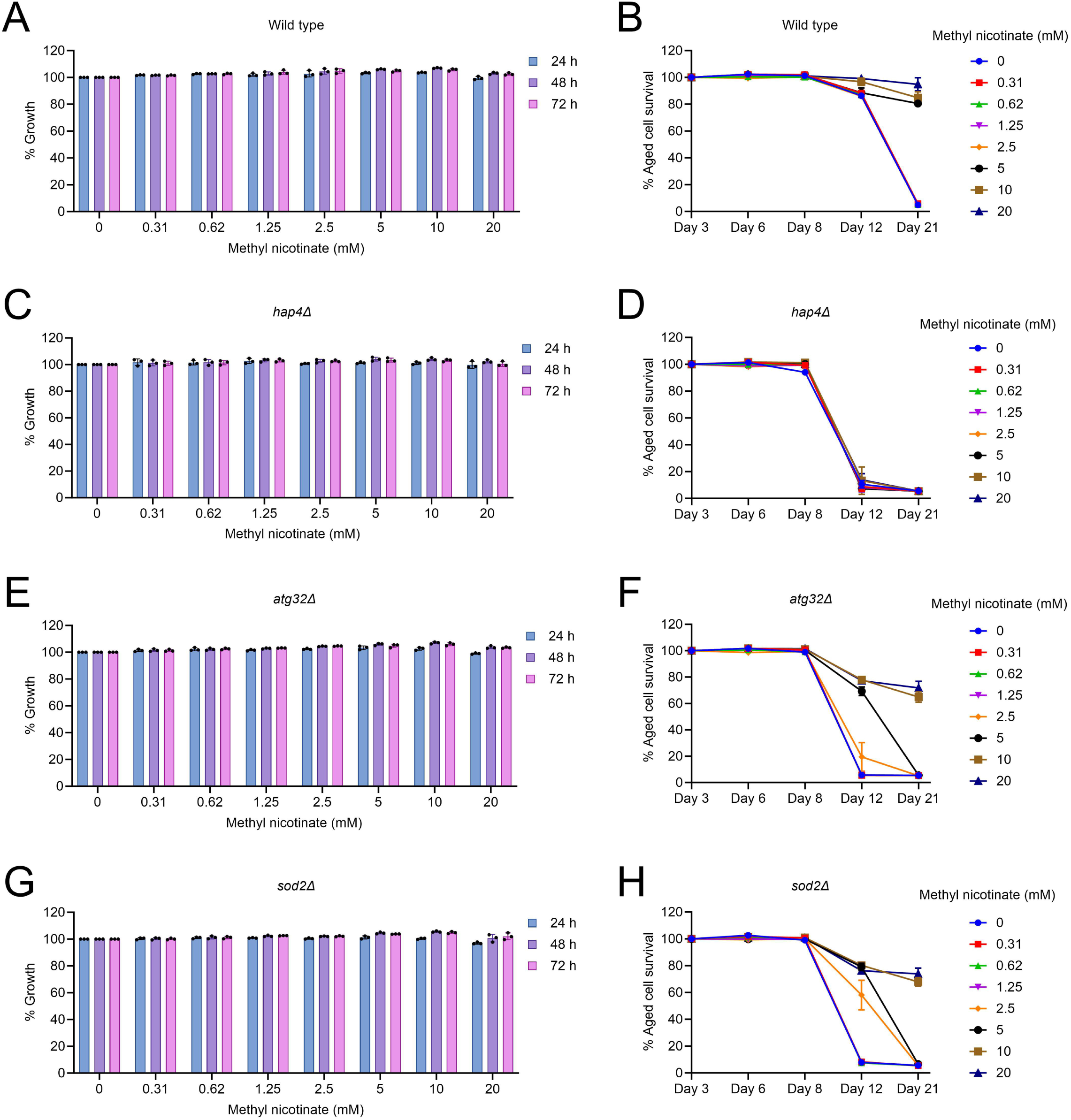
Methyl nicotinate (MN) induced lifespan extension via mitochondrial and stress resilience pathways. (A, C, E, G) Growth analysis of *Saccharomyces cerevisiae* prototrophic CEN.PK113- 7D wild type, *hap4Δ*, *atg32Δ* and *sod2Δ* cells in synthetic defined (SD) medium with indicated concentrations of MN supplementation assessed at 24, 48, and 72 hours using a 96-well plate. Data are presented as mean ± SD (n = 3). (B, D, F, H) Chronological lifespan (CLS) of wild type, *hap4Δ*, *atg32Δ* and *sod2Δ* was evaluated with indicated concentrations of MN supplementation. Survival was measured relative to day 3 outgrowth and plotted over time. Data are presented as mean ± SD (n = 3).

We also examined mutants exhibiting moderate mitochondrial dysfunction: *atg32Δ*, deficient in mitophagy ^38,39^, and *sod2Δ*, deficient in mitochondrial antioxidant defense ^40,41^. Both mutants displayed growth comparable to wild type but had slightly shorter lifespans (Figures 3E and 3G). MN supplementation at higher concentrations (10–20 mM) fully rescued lifespan in these strains, while lower concentrations were less effective (Figures 3F and 3H). These results suggest that MN can compensate for mild mitochondrial defects and oxidative stress, likely by enhancing mitochondrial function and stress response pathways.

Together, these data support a model in which MN extends lifespan via a mitochondria-dependent mechanism requiring *SNF1*-*HAP4*-mediated mitochondrial biogenesis and intact respiratory function, while also promoting mitochondrial quality control and oxidative stress resilience.

### Methyl nicotinate extends lifespan and improves mitochondrial function in human cells

To translate the conserved geroprotective effects of MN observed in yeast to mammalian systems, we next investigated its impact in human cells. Building on our findings that MN promotes mitochondrial biogenesis, quality control, and oxidative stress resilience in yeast, we assessed whether MN could also enhance cellular survival during chronological aging in human cell models.

We tested MN supplementation in human embryonic kidney-derived (HEK293) cells and human neuronal-like SH-SY5Y cells under chronological aging conditions ^28,42^. MN supplementation significantly improved the survival of both HEK293 (Figure 4A) and SH-SY5Y (Figure 4B) cells, indicating a conserved pro-longevity effect across species.

**Figure 4.**
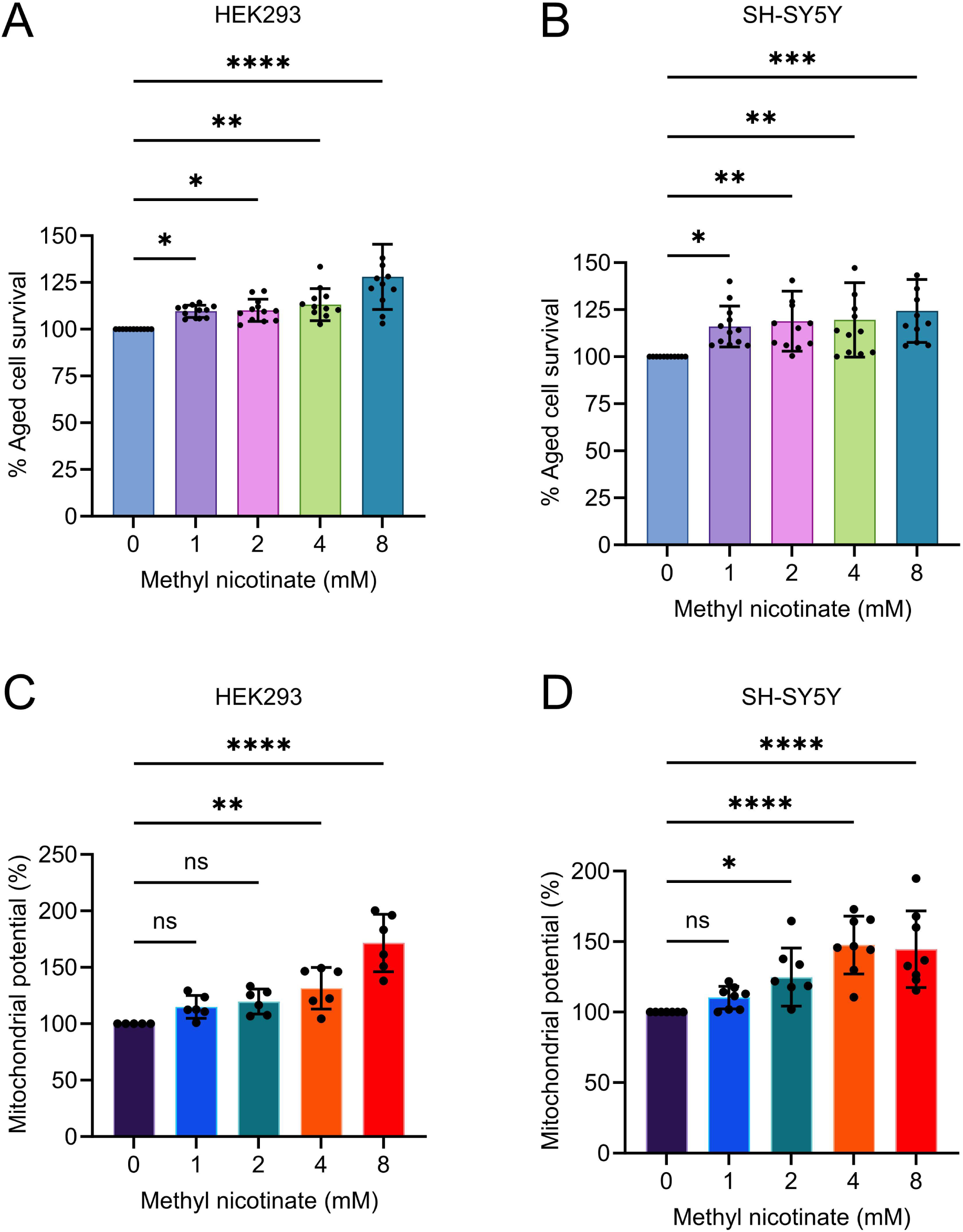
Methyl nicotinate (MN) enhances cellular survival and mitochondrial function in human cells. (A, B) Chronological aging assays in (A) human embryonic kidney-derived (HEK293) cells and (B) human neuronal-like SH-SY5Y cells supplemented with the indicated concentrations of MN. The survival of aged cells on Day 7 was using propidium iodide method. Cells survival of measured relative to the untreated control (100% survival). Data are represented as means ± SD (n=12). *P < 0.05, **P < 0.01, ***P < 0.001 and ****P < 0.0001 based on ordinary one-way ANOVA followed by Dunnett’s multiple comparisons test. two-way ANOVA followed by Dunnett’s multiple comparisons test. (C, D) Mitochondrial membrane potential was assessed using DiOC_6_ staining in HEK293 (C) and SH-SY5Y (D) cells supplemented with the indicated concentrations of MN. Fluorescence intensity was normalized to cell viability as determined by parallel CCK-8 assays. Data are presented as means ± SD (n = 6 for HEK293; n = 8 for SH- SY5Y). *P < 0.05, **P < 0.01, ****P < 0.0001, and ns = not significant, based on one- way ANOVA followed by Dunnett’s multiple comparisons test. (D-G) Cell survival analysis of HEK293 cells supplemented with varying concentrations of 4-MBA (D), 3-MBA (E) and 2-MBA (F) and BA (G).

To further explore the mechanism, we assessed mitochondrial function by measuring mitochondrial membrane potential—a key indicator of mitochondrial health^43^. MN supplementation enhanced mitochondrial membrane potential in both HEK293 and SH-SY5Y cells (Figures 4C and 4D), suggesting that MN promotes mitochondrial activity in human cells as well.

These findings demonstrate that MN not only extends lifespan in yeast via mitochondrial pathways but also enhances survival and mitochondrial function in human cells, underscoring its potential as a conserved geroprotective intervention.

### Methyl nicotinate interacts with the NAD⁺ biosynthetic pathway

Given the chemical nature of methyl nicotinate (MN) as a methyl ester of nicotinic acid (NA) ^29^, we hypothesized that MN may contribute to NAD⁺ biosynthesis and exert its lifespan-extending effects through this pathway. To investigate this, we assessed the growth response of key NAD⁺ biosynthetic deletion mutants, *npt1Δ*, *pnc1Δ*, and *bna6Δ* in the *Saccharomyces cerevisiae* BY4743 background grown in YPD medium. A schematic overview of the NAD⁺ biosynthetic network is shown in Figure 5A, highlighting that NA is generated from nicotinamide (NAM) via *PNC1*, converted to nicotinic acid mononucleotide (NaMN) via *NPT1*, and alternatively synthesized from tryptophan (TRP) via quinolinic acid (QA) in a *BNA6*-dependent *de novo* pathway ^34^.

**Figure 5.**
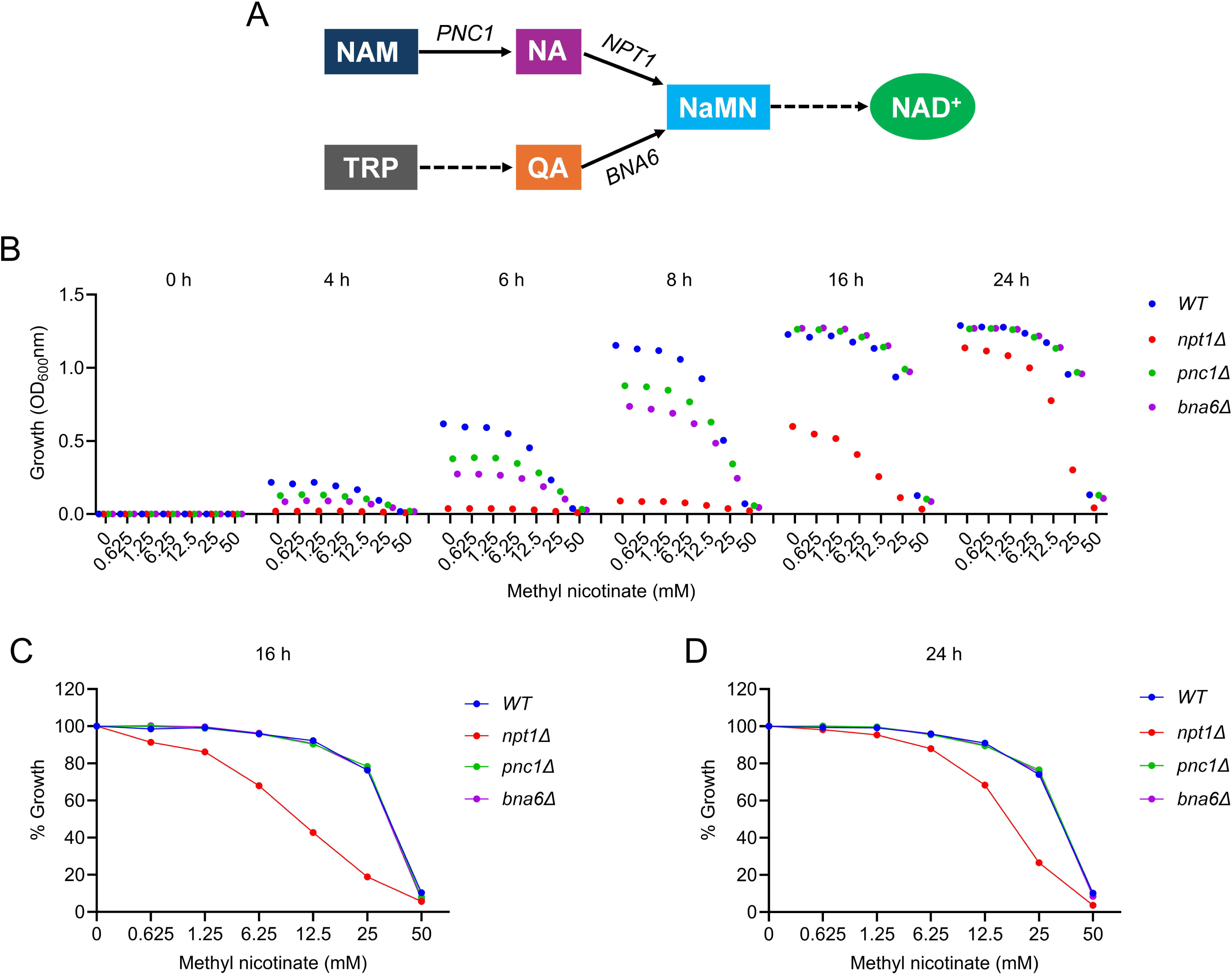
Methyl nicotinate (MN) interacts with the NAD⁺ biosynthetic pathway in *Saccharomyces cerevisiae.* (A) Schematic representation of the NAD⁺ biosynthesis pathways in yeast, including the salvage pathway from nicotinamide (NAM) via PNC1 and NPT1, and the de novo pathway from tryptophan via BNA6. (B) Growth kinetics of *Saccharomyces cerevisiae* auxotrophic BY4743 wild type (WT), *npt1Δ*, *pnc1Δ*, and *bna6Δ* cells in YPD medium supplemented with a 2-fold serial dilution of MN (0–50 mM), measured at 0, 4, 6, 8, 16, and 24 hours. (C, D) Quantification of MN-induced growth inhibition at 16 h (C) and 24 h (D) for all tested strains, shown as normalized growth relative to their respective untreated controls from panel B.

To assess MN’s interaction with these pathways, we performed a time-course growth assay over 0, 4, 6, 8, 16, and 24 hours. Cells were treated with a 2-fold dilution series of MN (0–50 mM) (Figure 5B). First, we characterized the basal growth dynamics of each strain (untreated). All deletion mutants displayed slower growth than wild type (Figure 5B) emphasizing the critical role of NAD⁺ biosynthesis in maintaining cellular growth. Among them, *npt1Δ* cells exhibited the most pronounced growth defect, entering exponential phase only after 16 hours (Figure 5B), while *pnc1Δ* and *bna6Δ* strains showed moderately reduced growth compared to wild type. These findings underscore the importance of intact NAD⁺ biosynthetic pathways for normal cell growth. However, whether this reduced growth results from NAD⁺ deficiency or accumulation of metabolic intermediates remains to be determined.

In wild type cells, MN inhibited growth at higher concentrations, indicating dose-dependent toxicity (Figures 5B-5D). Strikingly, the *npt1Δ* strain exhibited hypersensitivity to MN, showing pronounced growth inhibition even at lower concentrations (Figures 5B-5D). This suggests that *NPT1* is critical for processing MN—likely by converting MN-derived NA to NaMN, and that disruption of this step results in MN accumulation or failure to utilize MN for NAD⁺ biosynthesis. In contrast, *pnc1Δ* and *bna6Δ* strains displayed growth similar to wild type cells across all MN concentrations (Figures 5B-5D), indicating that MN does not depend strongly on NAM- to-NA conversion via *PNC1* or the de novo route through *BNA6* under these conditions.

To determine whether the hypersensitivity of the *npt1Δ* strain to MN is due to the accumulation of NA derived from MN, we profiled the growth response of all strains to nicotinamide (NAM). Since MN is structurally related to NA and could be metabolized into NA, we reasoned that NAM treatment would help reveal whether downstream accumulation of NA might underlie the observed toxicity in *npt1Δ* cells. NAM also exhibited dose-dependent toxicity in wild type cells, with marked growth inhibition evident as early as 6-8 hours post-treatment (Figure S1A). Notably, *npt1Δ* cells showed hypersensitivity to NAM, whereas the *pnc1Δ* strain did not, despite its inability to convert NAM to NA (Figures S1A-S1C). This pattern suggests that the accumulation of NA derived from NAM, rather than NAM itself, may be responsible for the growth inhibition. The lack of hypersensitivity in *pnc1Δ* cells further supports this conclusion, as these cells are unable to generate NA from NAM and would thus be protected from NA-mediated toxicity. Collectively, these findings implicate NA accumulation, particularly in the absence of *NPT1*, as a likely cause of growth defects observed under MN and NAM treatment.

To further investigate whether MN-induced toxicity in the *npt1Δ* strain results from the accumulation of NA, we next tested the growth response of all strains to exogenous NA. Although NA uptake in yeast is known to be limited due to poor membrane permeability ^34^, we observed that wild type cells remained largely unaffected by NA supplementation (Figure S2A-S2C), consistent with inefficient internalization. Interestingly, *npt1Δ* cells were markedly more sensitive to NA (Figure S2A-S2C), mirroring their hypersensitivity to both MN and NAM (Figures 5 and S2). In contrast, *pnc1Δ* and *bna6Δ* strains showed growth profiles comparable to wild type. These findings reinforce that the growth defects observed in *npt1Δ* cells are likely due to the intracellular accumulation of NA that cannot be further metabolized in the absence of *NPT1*. Together, this supports the conclusion that MN contributes to NAD⁺ biosynthesis via its conversion to NA and highlights *NPT1* as a critical node in mitigating potential metabolite toxicity.

### *IAH1* is a candidate esterase responsible for methyl nicotinate to nicotinic acid conversion in yeast

Given that MN is a methyl ester of NA, we hypothesized that its conversion to NA in *Saccharomyces cerevisiae* requires the activity of an esterase. To identify candidate enzymes potentially responsible for this hydrolysis step, we queried the Saccharomyces Genome Database (SGD) using the Gene Ontology (GO) term “hydrolase activity, acting on ester bonds.” This search returned five manually curated entries corresponding to the following genes: *LDH1*, *FSH1*, *DCR2*, *IAH1*, and *YPR147C*.

To determine whether any of these genes are involved in MN metabolism, we examined the growth profiles of the corresponding deletion strains in the presence of MN. Our rationale was that if the encoded hydrolase contributes to the conversion of MN to NA, its deletion would impair this process and result in increased sensitivity to MN due to accumulation of unmetabolized MN. Interestingly, among the five tested strains, the *iah1Δ* strain exhibited increased sensitivity to MN compared to wild type (Figures 6A-6C), indicating a possible role for *IAH1* in MN metabolism.

**Figure 6.**
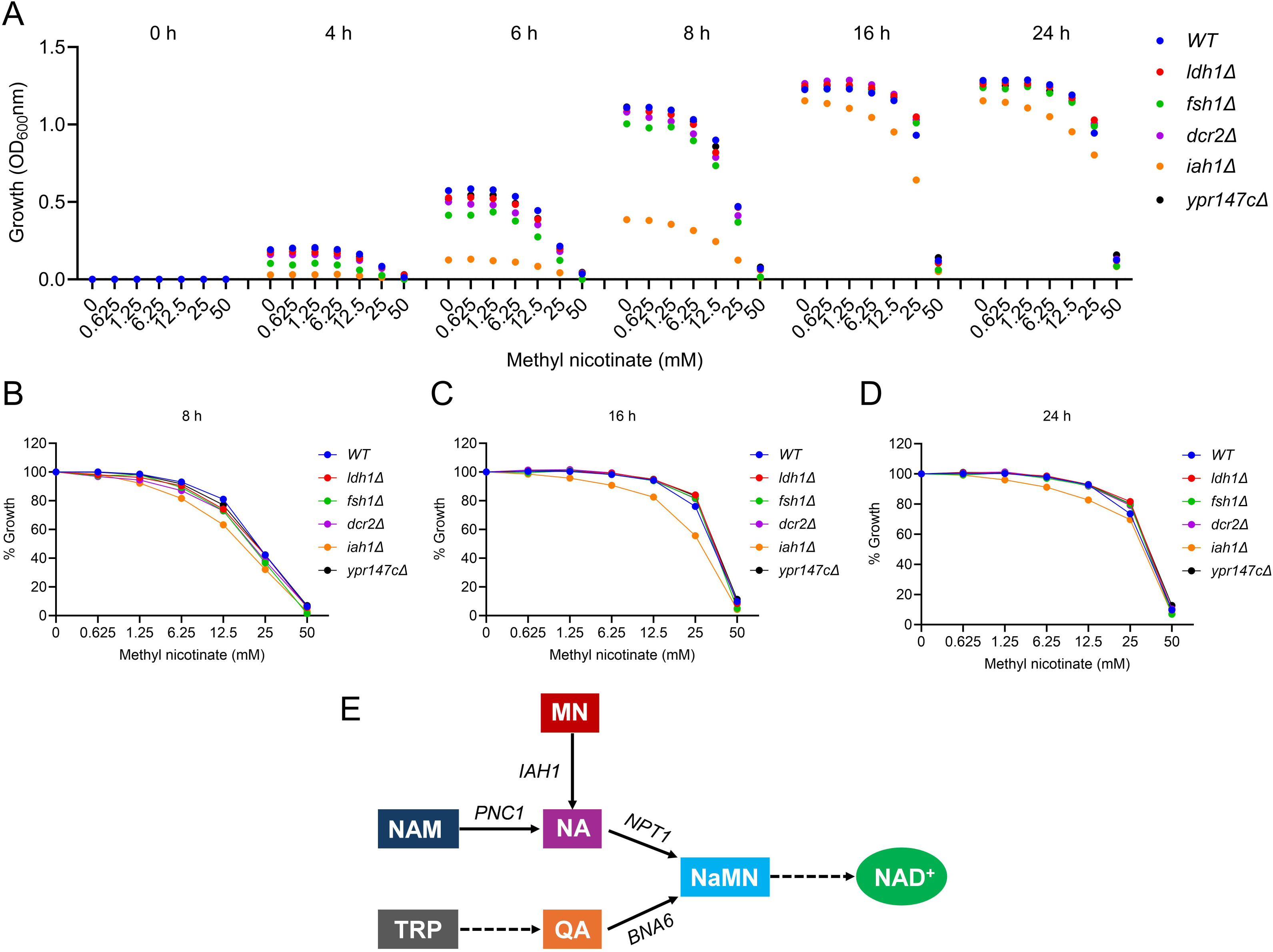
Screening of candidate esterases responsible for methyl nicotinate (MN) to nicotinic acid (NA) conversion in *Saccharomyces cerevisiae*. (A) Growth kinetics of *Saccharomyces cerevisiae* auxotrophic BY4743 wild type (WT), and esterase deletion strains (*ldh1Δ*, *fsh1Δ*, *dcr2Δ*, *iah1Δ*, and *ypr147cΔ*) grown in YPD medium supplemented with a 2-fold serial dilution of MN (0–50 mM), monitored over 24 hours. (B, C, D) Quantification of MN-induced growth inhibition at 8 h (B), 6 h (C) and 24 h (D) for all tested strains, shown as normalized growth relative to their respective untreated controls from panel A. (E) Proposed model of methyl nicotinate (MN) metabolism in *Saccharomyces cerevisiae*. MN enters the cell and is hydrolyzed by esterases *IAH1*, producing nicotinic acid (NA). NA then feeds into the NAD⁺ biosynthesis pathway via *NPT1*. This route functions in parallel with the salvage pathway from nicotinamide (NAM), which requires *PNC1*, and the *de novo* pathway from tryptophan, involving *BNA6*, together ensuring cellular NAD⁺ homeostasis.

To further investigate the role of *IAH1* in nicotinate metabolism, we profiled the growth of hydrolase deletion strains including *iah1Δ* in the presence of NAM and NA. While most deletion strains displayed growth patterns comparable to wild type cells, *iah1Δ* exhibited increased resistance to both NAM and NA, with the effect more prominent at early time points (Figures S3A-3C and S4A-4C). Given that both NAM and NA are toxic to wild type cells at higher concentrations, the resistance observed in *iah1Δ* may be due to a reduced conversion of MN to NA. This would result in lower intracellular NA levels, making the cells less susceptible to NA-derived toxicity.

*IAH1* encodes an esterase previously implicated in the hydrolysis of short-chain ester compounds ^44^, consistent with the structural characteristics of MN. While direct biochemical evidence is required to confirm its activity on MN, these results strongly suggest that *IAH1* may function as the hydrolase responsible for converting MN to NA in yeast. Interestingly, *iah1Δ* cells also showed slower basal growth (Figures 6, S3, S4), similar to the *npt1Δ* strain, which may reflect reduced intracellular NAD⁺ synthesis due to impaired MN conversion. While *NPT1*’s role in NAD⁺ biosynthesis and longevity regulation are well-established ^13,34,39,45^, our findings suggest that *IAH1* may play a previously unrecognized role in the same pathway. Interestingly, prior studies have also reported reduced lifespan in *iah1Δ* strains ^46^, further supporting a potential link between *IAH1* function, NAD⁺ metabolism, and aging.

## DISCUSSION

In this study, we identify methyl nicotinate (MN) as a novel compound that significantly extends the chronological lifespan (CLS) of *Saccharomyces cerevisiae* and enhances mitochondrial function and survival in human cells. Using a high-throughput chemical screen followed by detailed mechanistic dissection, we demonstrate that MN’s pro-longevity effect is mitochondria-dependent and mediated via the *SNF1-HAP4* pathway, implicating enhanced mitochondrial biogenesis, quality control, and oxidative stress resilience.

MN, a methyl ester of NA ^29^, is structurally positioned to modulate NAD⁺ biosynthesis, a central hub for cellular energy metabolism, stress response, and longevity ^14,17–20,33–35,47,48^. Consistent with this notion, MN extended lifespan in wild type yeast cells but failed to do so in mutants lacking functional mitochondria (*cox6Δ*) or the transcriptional activator of mitochondrial biogenesis, *HAP4* (*hap4Δ*). Importantly, MN could partially rescue lifespan in *snf1Δ* cells but not in *hap4Δ* or *cox6Δ* strains, suggesting that while *SNF1* enhances MN efficacy, mitochondrial integrity and *HAP4*-mediated biogenesis are essential for its function.

A key unresolved question was how MN exerts its effect intracellularly, given that methylated compounds like MN typically require hydrolysis by esterases to release the active carboxylic acid form (NA in this case). Our findings suggest that esterase activity is a critical determinant of MN processing and toxicity, particularly in the context of NAD⁺ metabolism.

We investigated this hypothesis by deleting *IAH1*, a known intracellular esterase, and observed that *iah1Δ* mutants were less sensitive to MN, supporting a role for *Iah1* in converting MN into NA. This functional evidence aligns with our observation that *npt1Δ* cells, which are unable to convert NA to NaMN, showed hypersensitivity to MN, NAM, and NA. Together, these data suggest that MN is metabolized into NA in an *IAH1*- dependent manner, and that in the absence of *NPT1*, NA accumulates, causing growth inhibition. Interestingly, *pnc1Δ* and *bna6Δ* strains showed no comparable sensitivity, reinforcing that the NA to NaMN step (via *NPT1*) is the critical bottleneck under MN treatment.

Thus, we propose a model in which MN enters the cell and is hydrolyzed by *IAH1* esterase, producing NA (Figure 6E). In wild type cells, NA is readily converted into NaMN by *NPT1*, feeding into NAD⁺ biosynthesis and supporting mitochondrial function and stress resistance. However, in *npt1Δ* mutants, this pathway is blocked, leading to toxic NA accumulation. Supporting this, both NAM and NA treatments recapitulated the MN toxicity pattern in *npt1Δ* mutants, but not in *pnc1Δ* or *bna6Δ* strains.

Interestingly, *IAH1* is conserved across species, including Homo sapiens, where it is annotated as isoamyl acetate hydrolyzing esterase 1 (putative) (Gene ID: 285148), also known as the isoamyl acetate-hydrolyzing esterase 1 homolog. It is a protein-coding gene predicted to be involved in lipid catabolic processes and identical protein binding, and may act upstream of or within gene expression regulation. Although the function of human *IAH1* is not fully understood, its predicted esterase activity raises the possibility that it may play a role in processing small esterified molecules. Whether human *IAH1* or a related esterase performs a similar MN bioactivation role remains to be determined. Future studies are warranted to explore its role in NAD⁺ metabolism, particularly in the context of MN conversion, mitochondrial regulation, and aging or metabolic diseases where NAD⁺ homeostasis is disrupted.

Finally, our cross-species analysis shows that MN also promotes survival and mitochondrial function in human cells, including both HEK293 and SH-SY5Y lines. MN improved mitochondrial membrane potential, indicating enhanced mitochondrial activity, which is consistent with the yeast data showing MN’s reliance on mitochondrial function and biogenesis.

Altogether, this study identifies MN as a novel lifespan-extending compound that enhances mitochondrial function and stress resistance in both yeast and human cells. Its effects are mitochondria-dependent and mediated through the *SNF1–HAP4* pathway, with *IAH1*-mediated bioactivation playing a key role in linking MN to NAD⁺ biosynthesis and cellular health.

## METHODS

### Yeast Strains, Media, and Culture Conditions

This study utilized both prototrophic *Saccharomyces cerevisiae* strain CEN.PK113-7D^49^ and the auxotrophic strain BY4743 (Euroscarf). Gene deletion mutants were constructed using a standard PCR-based gene disruption approach ^50^. Yeast strains were initially revived from glycerol stocks by streaking onto YPD agar plates (1% Bacto yeast extract, 2% Bacto peptone, 2% glucose, 2.5% Bacto agar) and incubated at 30°C for 2–3 days. Routine cultivation of CEN.PK113-7D was carried out in synthetic defined (SD) medium, composed of 6.7 g/L yeast nitrogen base with ammonium sulfate (without amino acids) and 2% glucose. In contrast, the auxotrophic BY4743 strains were maintained in rich YPD medium.

### Human Cell Culture and Growth Conditions

HEK293 (human embryonic kidney) and SH-SY5Y (human neuronal-like) cell lines were sourced from the American Type Culture Collection (ATCC, USA). Cells were maintained in high-glucose Dulbecco’s Modified Eagle Medium (DMEM; Cytiva, USA), supplemented with 10% heat-inactivated fetal bovine serum (FBS; Cytiva, USA) and 1% penicillin-streptomycin (Cytiva, USA). Cultures were incubated at 37°C in a humidified environment containing 5% CO_2_.

### Yeast Chronological Lifespan Assay

Chronological lifespan (CLS) analysis was carried out using prototrophic *Saccharomyces cerevisiae* strain CEN.PK113-7D cultured in synthetic defined (SD) medium composed of 6.7 g/L yeast nitrogen base with ammonium sulfate (without amino acids) and supplemented with 2% glucose ^8,24,31,51^. Overnight cultures grown at 30°C with constant agitation (220 rpm) were diluted to an initial OD600 of ∼0.2 in fresh SD medium to begin the assay. The assay was conducted in 96-well plates, each well containing 200 μL of culture, with compounds added in a two-fold serial dilution series; untreated wells served as controls. Cultures were incubated at 30°C, and growth was assessed at 24, 48, and 72 hours. To evaluate cell survival over time, an outgrowth-based method was employed. At the indicated time points, 2 μL of the aging culture was transferred into wells containing 200 μL of YPD medium in a fresh 96-well plate. Following a 24-hour incubation at 30°C without shaking, the OD600 was recorded using a microplate reader to estimate cell viability based on regrowth capacity.

### Yeast Growth Profiling Assay

To evaluate compound effects on yeast growth, a high-throughput growth assay was performed using 96-well microplates. The auxotrophic strain BY4743 was cultured in YPD medium and adjusted to an initial optical density (OD600) of ∼0.2. Cells were seeded into each well with a final volume of 200 µL, containing a series of two-fold serial dilutions of the test compounds. Plates were incubated at 30°C, and cell growth was monitored by measuring OD600 using a microplate reader. This setup enables efficient, parallel screening of compound-induced changes in yeast growth dynamics.

### Human Cells Chronological Lifespan Assay

To assess chronological lifespan (CLS), we monitored the survival of aging HEK293 (human embryonic kidney) and SH-SY5Y (human neuronal-like) cells using the previously established PICLS method ^28,42^. Briefly, cells were stained with propidium iodide (PI, 5 µg/ml) in culture medium and plated in 96-well plates, with or without the indicated compound treatments. Plates were incubated at 37°C in a humidified atmosphere containing 5% CO_2_ and shielded from light to prevent photobleaching. Cell viability was measured at defined aging intervals using a microplate reader set to 535 nm excitation and 617 nm emission wavelengths.

### Cell Viability Assay

Cell viability was assessed using the CCK-8 assay kit (Dojindo, Japan) following the manufacturer’s protocol. HEK293 and SH-SY5Y cells were plated in 96-well plates and exposed to treatments for 72 hours. After treatment, 10 µL of CCK-8 reagent was added to each well, and the plates were incubated at 37°C. Absorbance was then measured at 450 nm using a microplate reader to quantify cell viability.

### Mitochondrial Membrane Potential Measurement

Mitochondrial membrane potential was evaluated using the fluorescent dye 3,3’- dihexyloxacarbocyanine iodide (DiOC6) ^52^. HEK293 and SH-SY5Y cells were incubated with 50 nM DiOC6 in PBS for 30 minutes at 37°C. Following staining, fluorescence intensity was measured using a microplate reader or flow cytometer. To account for differences in cell number or viability, fluorescence values were normalized to cell viability data obtained from parallel CCK-8 assays performed on identically treated and seeded plates.

### Statistical Analysis

All experimental data were analyzed using GraphPad Prism version 10. Mean values, standard deviations, and significance levels were calculated, and data were visualized using appropriate graphical representations. Statistical comparisons were carried out using ordinary one-way ANOVA followed by post hoc multiple comparison tests. Significance levels are indicated in figures as follows: *P < 0.05, **P < 0.01, ***P < 0.001, and ****P < 0.0001; results not reaching significance are labeled as ’ns’ (not significant). A P value below 0.05 was considered statistically significant.

### Data availability

Further information and requests for resources and reagents should be directed to and will be fulfilled by the Lead Contact, Dr. Mohammad Alfatah.

## ACKNOWLEDGMENTS

We thank Prof. Brian Kennedy (Director, Healthy Longevity Translational Research Programme, NUS Medicine, Singapore) and Prof. Koh Woon Puay (Healthy Longevity Translational Research Programme, NUS Medicine, Singapore) for their generous support of this research. We are also grateful to Dr. Wong Siew Peng Esther (Department of Physiology, NUS, Singapore) for her valuable support. Special thanks to Mr. Koh Ting Wei Kelvin (Healthy Longevity Translational Research Programme, NUS Medicine, Singapore) for his assistance in laboratory management. This work was supported by the Academic Health Programme (AHP) Fund 2024, National University Health System (NUHS), Singapore.

## AUTHOR CONTRIBUTIONS STATEMENT

Gao Mingtong: Investigation and Formal analysis

Mohammad Alfatah: Conceiving of the project, Writing and Funding Acquisition

## SUPPLEMENTARY FIGURE LEGENDS

**Figure S1.**
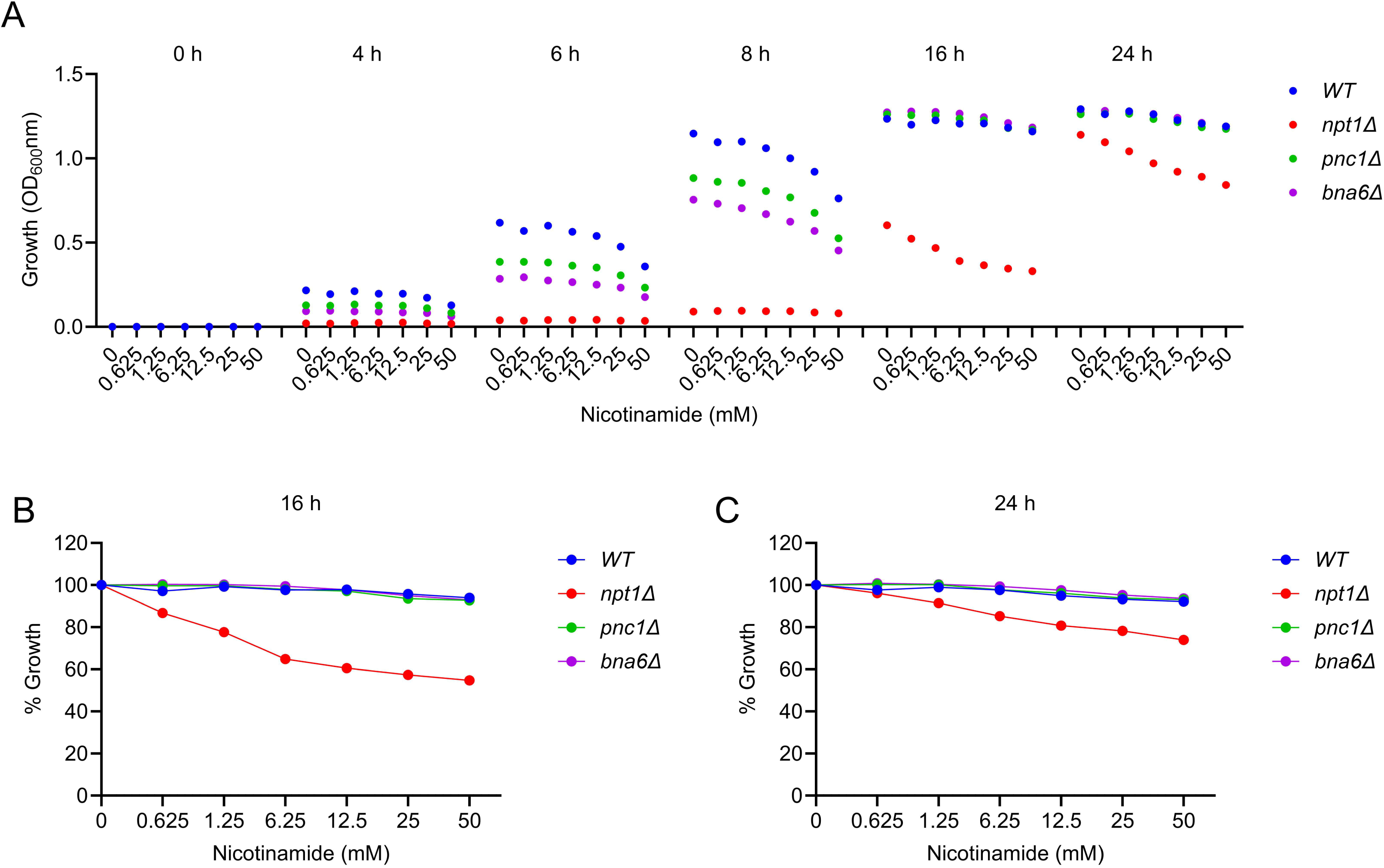
Growth profiling of NAD⁺ biosynthetic pathway mutants in response to nicotinamide (NAM). (A) Growth kinetics of *Saccharomyces cerevisiae* auxotrophic BY4743 wild type (WT), *npt1Δ*, *pnc1Δ*, and *bna6Δ* cells in YPD medium supplemented with a 2-fold serial dilution of NAM (0–50 mM), measured at 0, 4, 6, 8, 16, and 24 hours. (B, C) Quantification of NAM-induced growth inhibition at 16 h (B) and 24 h (C) for all tested strains, shown as normalized growth relative to their respective untreated controls from panel A.

**Figure S2.**
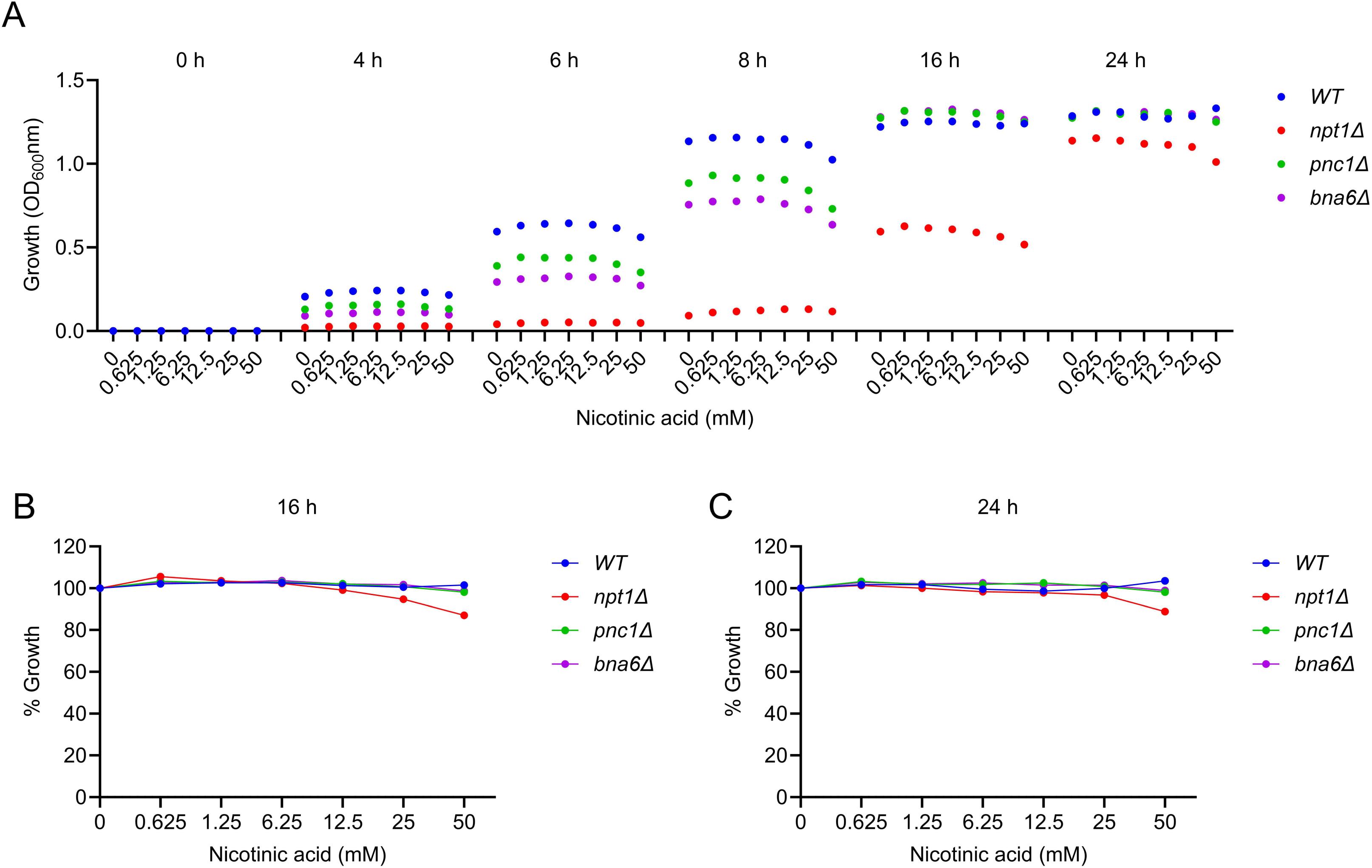
Growth profiling of NAD⁺ biosynthetic pathway mutants in response to nicotinic acid (NA). (A) Growth kinetics of *Saccharomyces cerevisiae* auxotrophic BY4743 wild type (WT), *npt1Δ*, *pnc1Δ*, and *bna6Δ* cells in YPD medium supplemented with a 2-fold serial dilution of NA (0–50 mM), measured at 0, 4, 6, 8, 16, and 24 hours. (B, C) Quantification of NA-induced growth inhibition at 16 h (B) and 24 h (C) for all tested strains, shown as normalized growth relative to their respective untreated controls from panel A.

**Figure S3.**
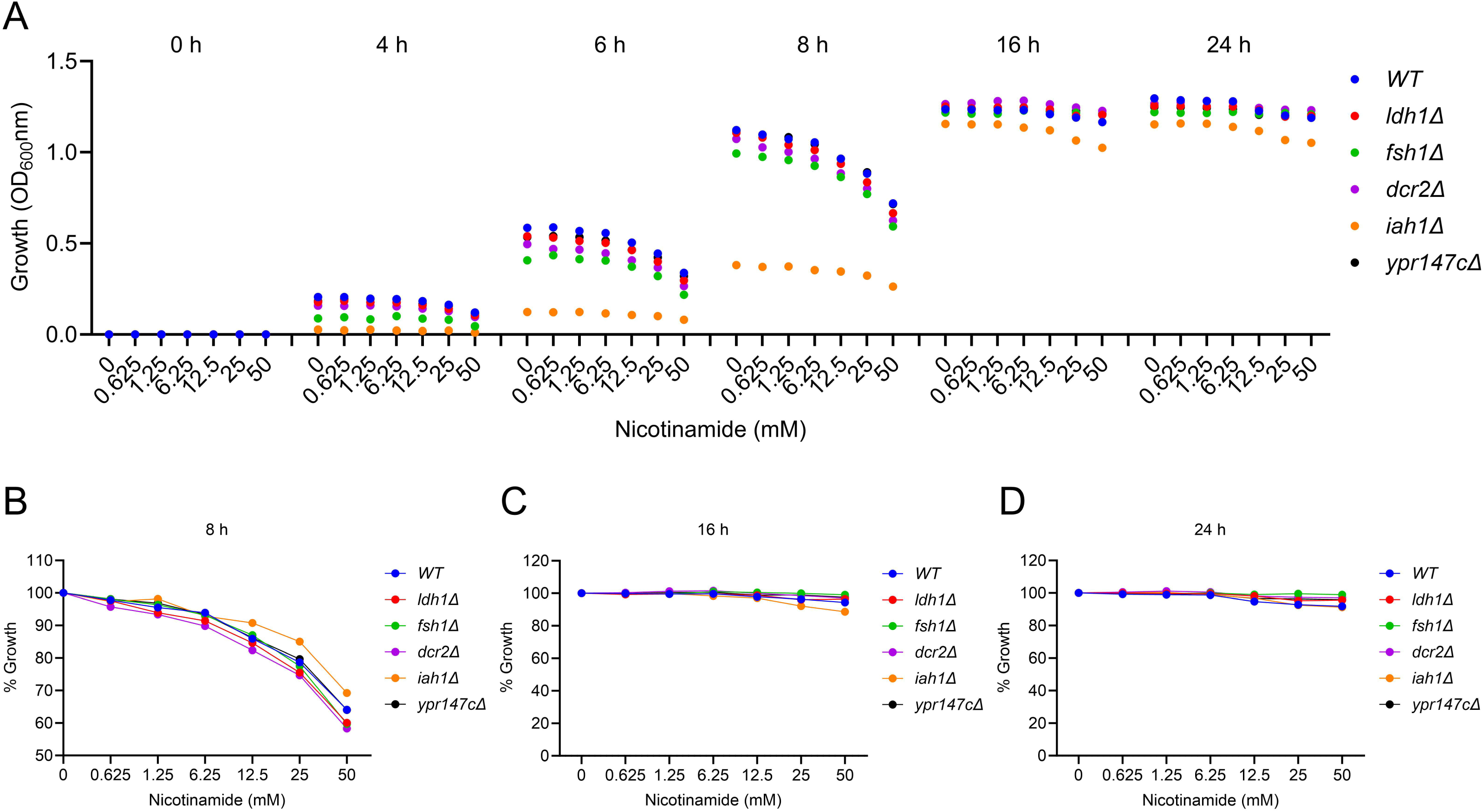
Growth profiling of candidate esterases mutants in response to nicotinamide (NAM). (A) Growth kinetics of *Saccharomyces cerevisiae* auxotrophic BY4743 wild type (WT), and esterase deletion strains (*ldh1Δ*, *fsh1Δ*, *dcr2Δ*, *iah1Δ*, and *ypr147cΔ*) grown in YPD medium supplemented with a 2-fold serial dilution of NAM (0–50 mM), monitored over 24 hours. (B, C, D) Quantification of NAM-induced growth inhibition at 8 h (B), 6 h (C) and 24 h (D) for all tested strains, shown as normalized growth relative to their respective untreated controls from panel A.

**Figure S4.**
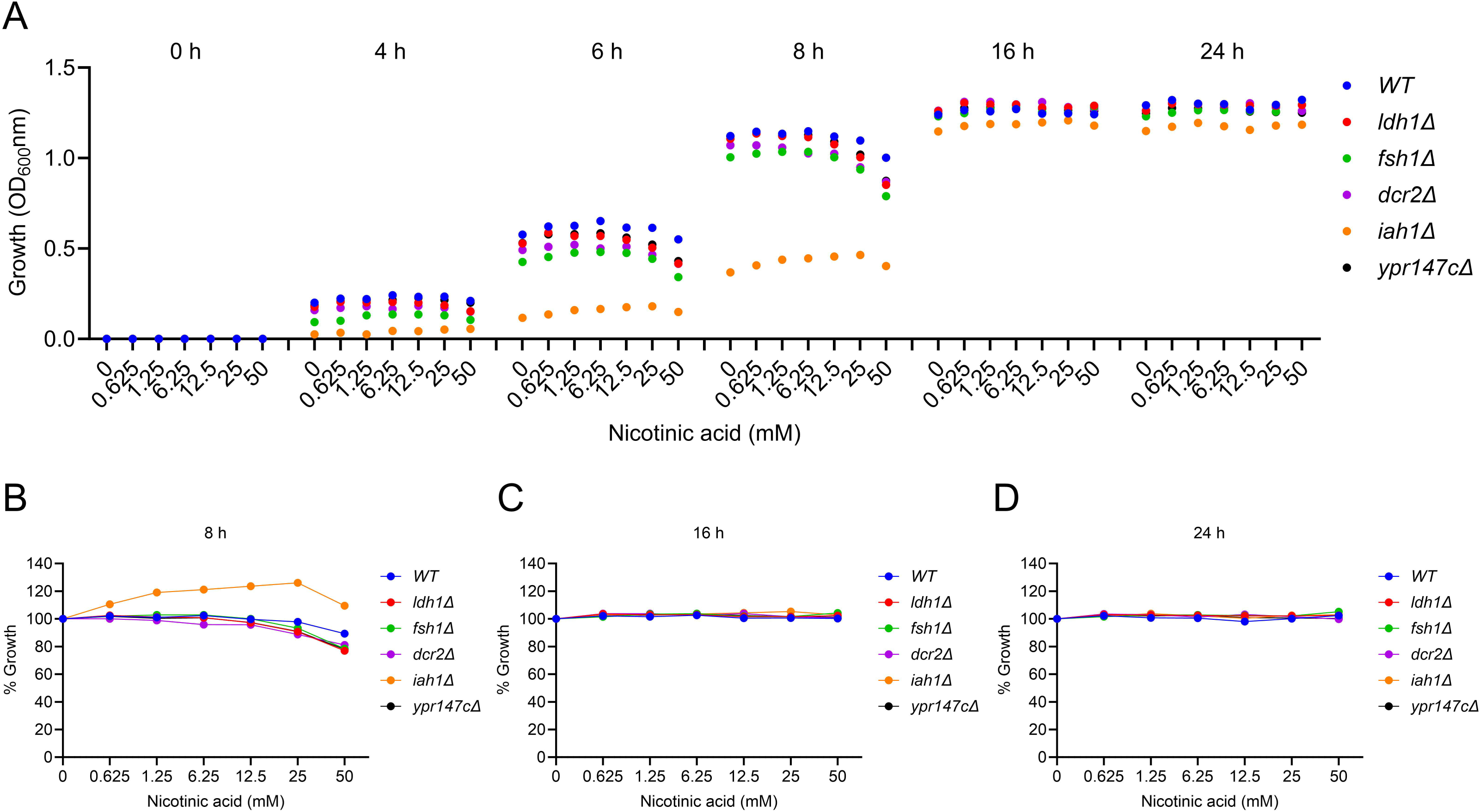
Growth profiling of candidate esterases mutants in response to nicotinic acid (NA). (A) Growth kinetics of *Saccharomyces cerevisiae* auxotrophic BY4743 wild type (WT), and esterase deletion strains (*ldh1Δ*, *fsh1Δ*, *dcr2Δ*, *iah1Δ*, and *ypr147cΔ*) grown in YPD medium supplemented with a 2-fold serial dilution of NA (0–50 mM), monitored over 24 hours. (B, C, D) Quantification of NA-induced growth inhibition at 8 h (B), 6 h (C) and 24 h (D) for all tested strains, shown as normalized growth relative to their respective untreated controls from panel A.

